# *Candida auris* can acquire antifungal resistance without selection pressure

**DOI:** 10.64898/2026.01.07.695227

**Authors:** Trinh Phan-Canh, Duc-Minh Nguyen-Le, Manju Chauhan, Phuc-Loi Luu, Anuradha Chowdhary, Neeraj Chauhan, Karl Kuchler

## Abstract

The human fungal pathogen *Candidozyma auris (formerly Candida auris*) can cause prolonged infection outbreaks with high mortality rates in healthcare settings. Treatment failures of patients arise not only from antifungal drug resistance, but also from intra-species variability in pathogenicity as well as induced hypermutation events in response to clinical therapy. Whole genome sequencing was used to identify genetic mutations using the CDC mycoSNPs pipeline. Antifungal susceptibility testing was performed based on Clinical and Laboratory Standards Institute (CLSI) methods. We report here that the interlaboratory exchange of *C. auris* clinical isolates dried on sterile filter paper, resulted in the emergence of at least three distinct morphotypes following reconstitution. These distinct morphotypes exhibited differences in drug resistance and morphogenesis, linked to the accumulation of mutations in genes associated with azole and echinocandin resistance. Using whole genome sequencing, we identified several variants in *TAC1B*, *MRR1* and *FKS2* that correlate with altered drug susceptibilities. Experiments recapitulating filter paper shipment conditions revealed genetic and epigenetic changes, explaining the morphogenetic switching and altered azole resistance. Our findings demonstrate that *C. auris* can acquire mutations affecting drug resistance traits even in the absence of antifungal exposure, raising concerns about shipment preparation procedures across mycology laboratories. The results are of broad relevance for the medical mycology community, as they call for standardized protocols for exchanging clinical strains, but also experiments to verify phenotypic traits between laboratories.

**Importance:** Pathogenesis and antifungal drug resistance traits of *Candida auris* vary widely across clinical strains and are often attributed to elevated mutation rates. In fungal pathogen research, clinical strains are commonly exchanged between laboratories by transfer on filter paper, a convenient and widely used practice. However, in the case of a *C. auris* clinical strain received from a collaborating laboratory, we identified multiple acquired mutations. These genetic alterations caused marked changes in morphogenesis and antifungal susceptibility, demonstrating that resistance in *C. auris* can arise without antifungal selection pressure. Our findings highlight the potential for genetic and phenotypic diversification during routine strain handling and underscore the need for standardized protocols for exchanging clinical *C. auris* strains across mycology laboratories.

## INTRODUCTION

*Candida auris* is an emerging human fungal pathogen characterized by unprecedented pan-antifungal multidrug resistance traits. Mortality rates of disseminated infections sometimes exceed 60% [1–3]. *C. auris* has a remarkable capacity for adaptive microevolution, often resulting in the emergence of new pathogenic traits, including altered morphogenesis, virulence [4] as well as pan-antifungal resistance [5–9]. Recent studies hint that immune defense or selection pressure from antifungal therapy can drive the emergence of drug resistance by accelerated mutations in numerous genes [7]. These findings support the notion that the skin-tropic *C. auris* pathogen can quickly diversify and adapt resistance levels and virulence traits during transmission in healthcare environments or when systemically disseminated in colonized individuals.

*C. auris* exhibits high resistance rates to all major antifungal classes including azoles (up to 90%), amphotericin B (up to 50%) and echinocandins (2-8%) [3]. Azole resistance is commonly associated with mutations in the gene encoding the *Erg11* lanosterol 14α-demethylase (Y132F, K143R, and VF125AL), increasing fluconazole minimal inhibitory concentrations (MICs) by ∼8-16-fold [10]. Further, gain-of-function (GOF) mutations such as A640V, A657V or ΔF862N866 in Tac1B upregulate the ATP-binding cassette (ABC) transporter Cdr1, driving up to ∼16-fold azole MIC increases [11–13]. The major facilitator superfamily (MFS) transporter Mdr1 also contributes, controlled by the transcription factor Mrr1 [14]. The most common Mrr1 GOF variant N647T elevates fluconazole MICs by ∼4-fold [10,14]. These partially redundant mechanisms explain the pronounced azole resistance of *C. auris*. Besides ergosterol, sphingolipids may further modulate azole and polyene responses [15,16]. Indeed, loss of the phosphatidyl inositol-transfer protein Pdr16 lowers azole MICs by ∼4-fold, whereas increased *PDR16* dosage can confer resistance to both azoles and amphotericin B (AMB) in *C. auris* [15,17]. Echinocandin resistance is predominantly associated with hotspot-1 substitutions in *FKS1* (S639F/P/Y) [2,18–20], while the role of *C. auris FKS2* remains unclear. Increasing evidence suggests the pronounced AMB resistance in *C. auris* involves multiple intrinsic pathways, including sphingolipid metabolism [9], CO_2_-sensing [21], mitochondrial function [21,22] and chromatin regulation [23], whereas adaptive AMB resistance rarely arises from mutations in ergosterol biosynthesis [5,24].

During antifungal therapy, drug-resistant *C. auris* isolates are increasingly recovered from patients [24,25], underscoring the roles of selection pressure and rapid microevolution [4,7]. In clinical settings, *C. auris* colonizes nutrient-lacking abiotic surfaces of healthcare devices, environmental fomites, as well as biotic surfaces such as human skin where it is frequently exposed to disinfectants [26,27]. Despite these constraints, *C. auris* can cause prolonged hospital outbreaks lasting 1-3 years [28–31], raising pivotal questions about its remarkable stress tolerance. Our recent work shows that *C. auris* can acquire antifungal resistance traits and stress resilience via epigenetic white-brown switching as well as *de novo* mutations [8,32], highlighting its pronounced phenotypic plasticity. The remarkable intra-species variations may be linked to climate change that can drive adaptive evolution with unprecedented dynamics and even facilitate switch in host range [33]. Moreover, certain epigenetic mechanisms can accelerate mutational rates [32]. These observations prompt the question as to whether *C. auris* can acquire drug-resistance traits even in the absence of direct antifungal selection.

Here, we show that a single *C. auris* clinical isolate may produce a mixed population of progeny exhibiting distinct morphologies, drug resistance as well as virulent traits. While our data show that this can occur within a given laboratory on filter paper, we cannot exclude that mutational changes arise during long-range transport or storage during transit. Our experiments, which simulate transfer conditions of strains dried on sterile filter paper, reveal a rapid accumulation of mutations linked to antifungal susceptibility and morphogenesis. These findings highlight the extensive intraspecies variations in *C. auris* and underscore the need for caution when transferring strains between mycology laboratories.

## RESULTS

### A mixture of three distinct phenotypes emerging from a pure clinical isolate

The clinical isolate CK52/P/13 was originally obtained from a collaborating laboratory, preserved on filter paper, and immediately reconstituted on YPD agar upon receipt, followed by conservation as a frozen stock in 20% glycerol at −80 °C **(Fig. 1A)**. Notably, after a few days on YPD agar at 37 °C but not at 30 °C, three distinct colony morphologies appeared yielding smooth (Smo), small (Sma), and wrinkled (Wri) **(Fig. 1A)** colonies. Morphological assessments on YPD agar at different temperatures showed that Smo colonies consisted of homogeneous yeast-form cells, while the Sma ones had aggregated cells with occasionally elongated forms **(Fig. 1B)**. Strikingly, Wri colonies growing at 37°C displayed a high proportion of abnormally elongated cells with massive vacuoles, as well as some with pseudohyphal morphologies. Interestingly, the phenotypes were much less pronounced at 30D°C or 42D°C. Antifungal dose-response assays using four classes of drugs revealed significant MIC variations of all morphotypes **(Fig. 1C)**. Both smooth and small morphotypes showed a 4-fold increase in MICs for caspofungin (CAS), while Sma and Wri colonies displayed an 8-fold MIC increase for voriconazole (VRC). Apparent fitness differences of colony morphotypes growing on YPD agar **(Fig. 1A-B)** implied possible trade-offs associated with drug resistance emergence.

**Figure 1.**
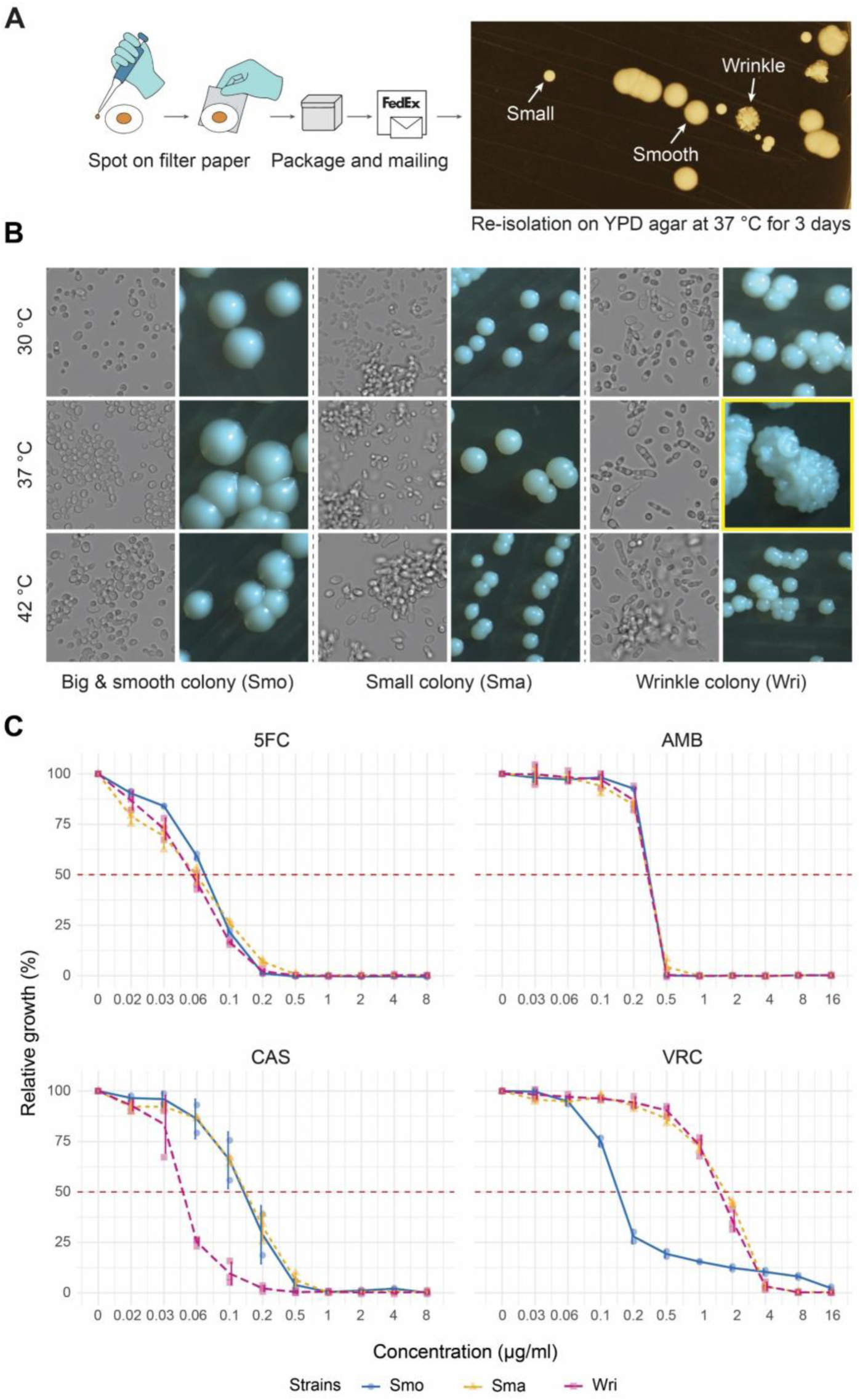
A mixed population of *C. auris* cells with distinct morphologies. **A.** Strain reconstitution from dried filter paper after inter-laboratory exchange. The YPD plate showed three distinct colony morphologies emerging from a single pure isolate after reconstitution. **B.** Cell (magnification 60x) and colony morphology (magnification 18x) of *C. auris* morphotypes growing at different temperatures on YPD agar for 3 days. **C.** Inhibitory dose response assays of *C. auris* morphotypes for flucytosine (5FC), amphotericin B (AMB), caspofungin (CAS) and voriconazole (VRC); n=2-3.

### Whole genome sequencing explains antifungal susceptibility and morphology variations

To test whether genomic mutations can account for underlying causes of morphotype alterations, we subjected colony morphotypes to whole-genome sequencing **(Fig. 2A)**. Remarkably, sequencing revealed multiple mutations in genes unique to the Sma morphotype, including B9J08_003129 (p.Y127fs), a xenosiderophore transporter *SIT1.4* (B9J08_001519 – p.F500fs), a histone H3K79 methyltransferase *DOT1* (B9J08_000044 – p.V94A) and a poly(A) polymerase *PAP1* (B9J08_001453 – p.T169R). *PAP1* is essential for viability, biofilm formation, and virulence in *C. albicans* [34,35], suggesting that a mutation in *C. auris* may impair fitness. Further, the Sma and Wri morphotypes shared many mutations, including changes in genes involved in cell separation and development **(Fig. 2B)** such as the regulator repressor *NRG1* (B9J08_005429), *TRA1* (B9J08_005406), *POP7* (B9J08_004414), *VAN1* (B9J08_003572), *KIN3*(B9J08_002993), and the *WOR2* transcription factor (B9J08_002136). Of note, *KIN3* is also linked to aggregation in *C. auris* [4], while *VAN1* appears essential for immune evasion, since Van1 maintains the outer cell wall mannan layer [4,36]. Additionally, Sma and Wri cells harbored frameshift mutations in the cAMP-dependent protein kinase *TPK1* (B9J08_004030 – p.L159fs) and the C_2_H_2_ zinc finger transcription factor *STP1.2* (B9J08_001104 – p.E378fs) potentially involved in morphogenesis [37]. These data may explain, at least in part, both altered morphogenesis and reduced fitness of these strains [38,39].

**Figure 2.**
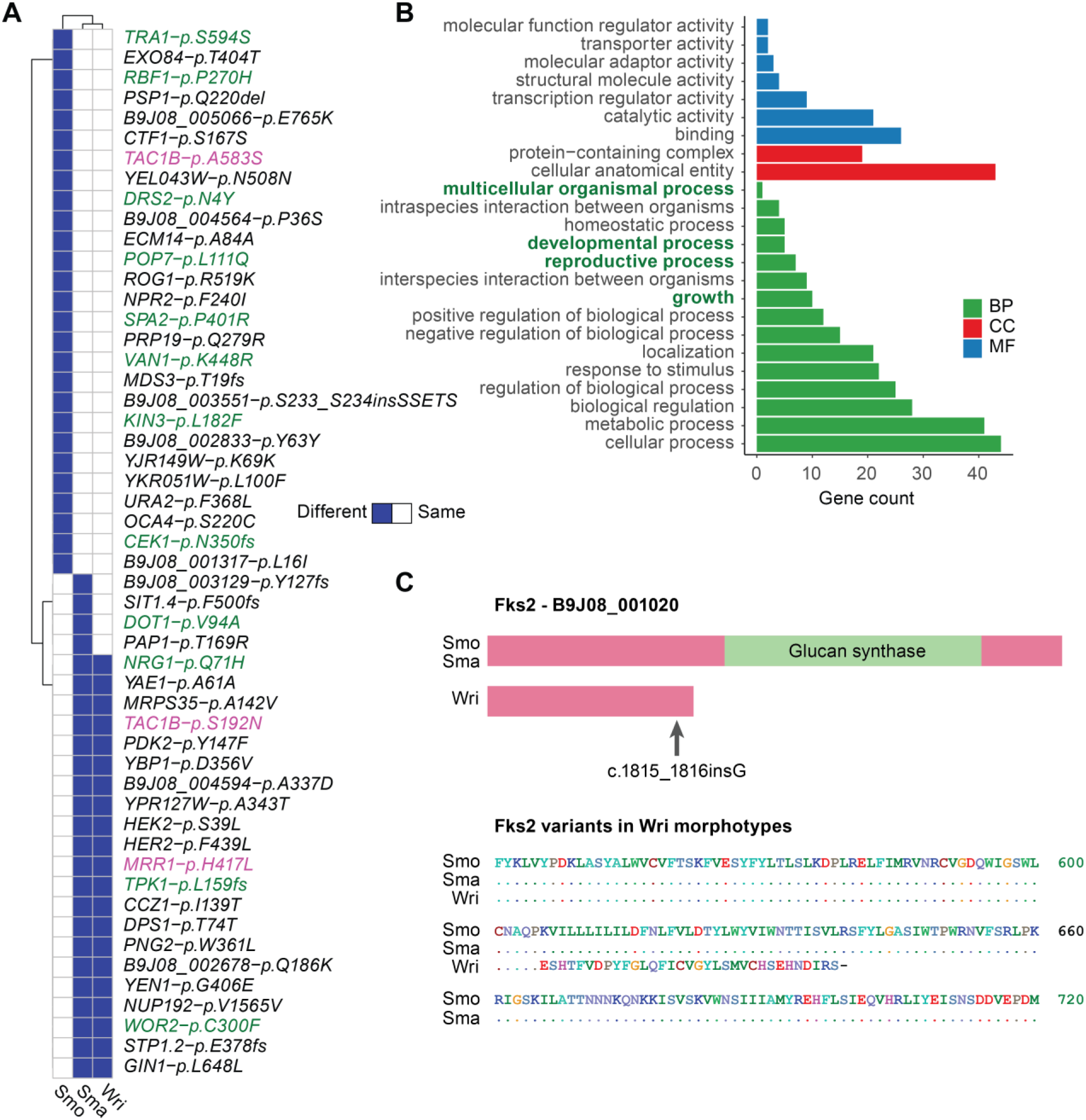
Genetic mutations associated with antifungal susceptibility and morphogenesis. **A.** Whole genome sequencing of a clinical isolate revealed numerous polymorphisms in genes linked to drug resistance, morphology alterations and fitness when compared to the standard strain B8441. Green color indicated genes related to biological processes shown in B. Pink color indicated genes associated with drug resistance. **B.** groupGO analysis of genes with mutations revealed GO terms related to altered morphological traits. **C.** A new frameshift mutation in *FKS2* producing a truncated glucan synthase explains enhanced caspofungin sensitivity.

Regarding antifungal resistance, both Sma and Wri colonies carried mutations in the master transcription factors *TAC1B* (S192N) and *MRR1* (H417L) **(Fig. 2A)**, regulating the efflux transporters Cdr1 and Mdr1 for azoles in *C. auris,* respectively, [14,40], as well as in other *Candida* spp [13,41,42]. The Tac1b^S192N^ variant, though less common in clinical strains [40], has been associated with a 4-fold MIC increase for fluconazole, most likely by activating the Cdr1-mediated ABC transporter efflux system [40]. The newly identified Mrr1^H417L^ variant may contribute additional azole resistance mechanisms by driving efflux permeases or controlling uptake [14].

Interestingly, Smo and Sma morphotypes displayed a marked CAS hypertolerance. While mutations could explain this phenotype, it remains unclear from our variant calling data **(Fig. 2A)**. Hence, we inspected the *FKS1/2* loci using the Integrative Genomics Viewer (IGV) tool [43]. Indeed, we found an as yet unknown mutation, *FKS2* (B9J08_001020), showing an allele frequency of approximately 50% **(Fig. S1)**, implying that this *FKS2* mutant variant can contribute to CAS resistance. Sanger sequencing confirmed a guanine insertion between nucleotide position 1815 and 1816 of *FKS2*, leading to frameshift at Gly606 and introducing a premature stop codon 33 amino acids downstream (p.Gly606fs*33) **(Fig. 2C)**. This result suggests that loss-of-function mutation in *FKS2* can significantly reduce the MIC for caspofungin by around 4-fold.

### *S*hipment on filter paper alters antifungal susceptibility phenotypes

To test whether morphotype diversities can emerge during or after strain shipment, we replicated the transfer conditions shown in **Fig. 1A**, adding a control in which the filter paper was kept moist with 50 µl of sterile water in a sealed microcentrifuge tube. After two weeks, *C. auris* cells were recovered and recultivated on YPD agar **(Fig. 3A)**. Abnormal colonies emerged from all morphotypes, though at a lower frequency when the filter paper was kept moist. The Sma and Wri morphotypes exhibited a higher diversification, possibly due to mutations in genes associated with DNA damage response and chromatin organization such as *DOT1, TRA1*, and *POP7* [39,44–46].

**Figure 3.**
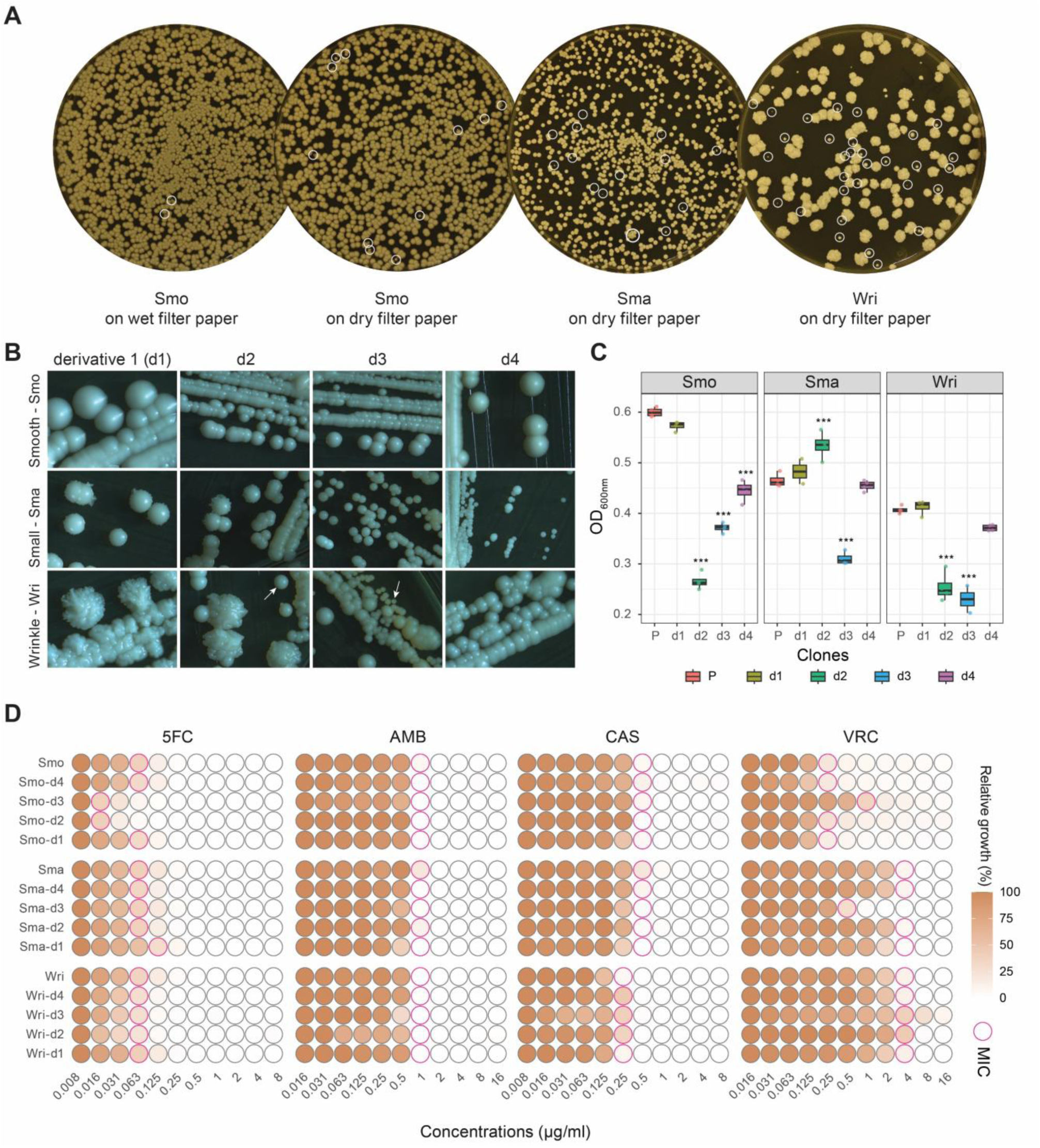
Drying of *C. auris* isolates on filter paper promotes emergence of new morphotypes. **A.** *C. auris* cell suspensions (∼ 10^9^ cells/ml) were spotted onto sterile Whatman filter paper. Wet filter papers were sealed in microcentrifuge tubes, while dried filters were folded in sterile aluminum foil. Preserved wet and dry colonies were re-isolated on YPD agar after two weeks. White circles indicate new morphotypes. **B.** Four colonies from each parental strain were re-streaked on YPD agar at 37°C and preserved for further experiments. **C.** Biomass of different derivatives in (B) growth in RPMI-MOPS pH7 for 24 hours at 37°C measured by recording OD_600nm_. **D.** Dose response MIC assay of derivatives collected in (B). MIC wells indicated threshold 50% growth inhibition compared to no-drug control well for all drugs, except a threshold 90% chosen for AMB.

We then inspected phenotypes of four dried derivative colonies from each morphotype on YPD at 37D°C reconstituted from filter papers. Notably, derivatives Wri-d2 and Wri-d3 from the wrinkled strain were able to switch between morphotypes **(Fig. 3B)**. For instance, Wri-d2 reversibly changed between Wri and Smo colonies, and Wri-d3 between large and small colony forms, implying epigenetic causes of switching **(Fig. 3B)**. This may explain, at least in part, the high rate of morphotype diversification in the Wri strain **(Fig. 3A)**. Furthermore, Wri derivatives showed significantly altered fitness traits in RPMI medium, but surprisingly only modest changes in drug susceptibility **(Fig. 3C)**. The Smo-d2 and Smo-d3 derivatives from the Smo colony exhibited hypersensitivity to 5FC, with a 4-fold MIC decrease for 5FC **(Fig. 3D)**. Meanwhile, Smo-d3 showed a 4-fold MIC increase to VRC. By contrast, Sma-d3 displayed reduced VRC MIC. These observations support the notion of fitness trade-offs, as the azole-resistant Smo-d3 cells showed impaired growth in RPMI. Similarly, Sma and Wri morphotypes were azole-resistant, with reduced fitness in the MIC-testing medium RPMI **(Fig. 3C)**.

### *De novo* mutations rapidly accumulate during preservation on filter paper

To assess whether mutagenesis occurred during preservation on filter paper or during shipment, we performed WGS of all original isolates and the derived strains **(Fig. 4A-C)**. Notably, DNA sequence changes were detected in all recovered clones, indicating mutagenic events mounting during preservation on filter paper regardless of moisture conditions. This observation suggested that *C. auris* may evolve rapidly when its colonies persist on surfaces in healthcare facilities.

**Figure 4.**
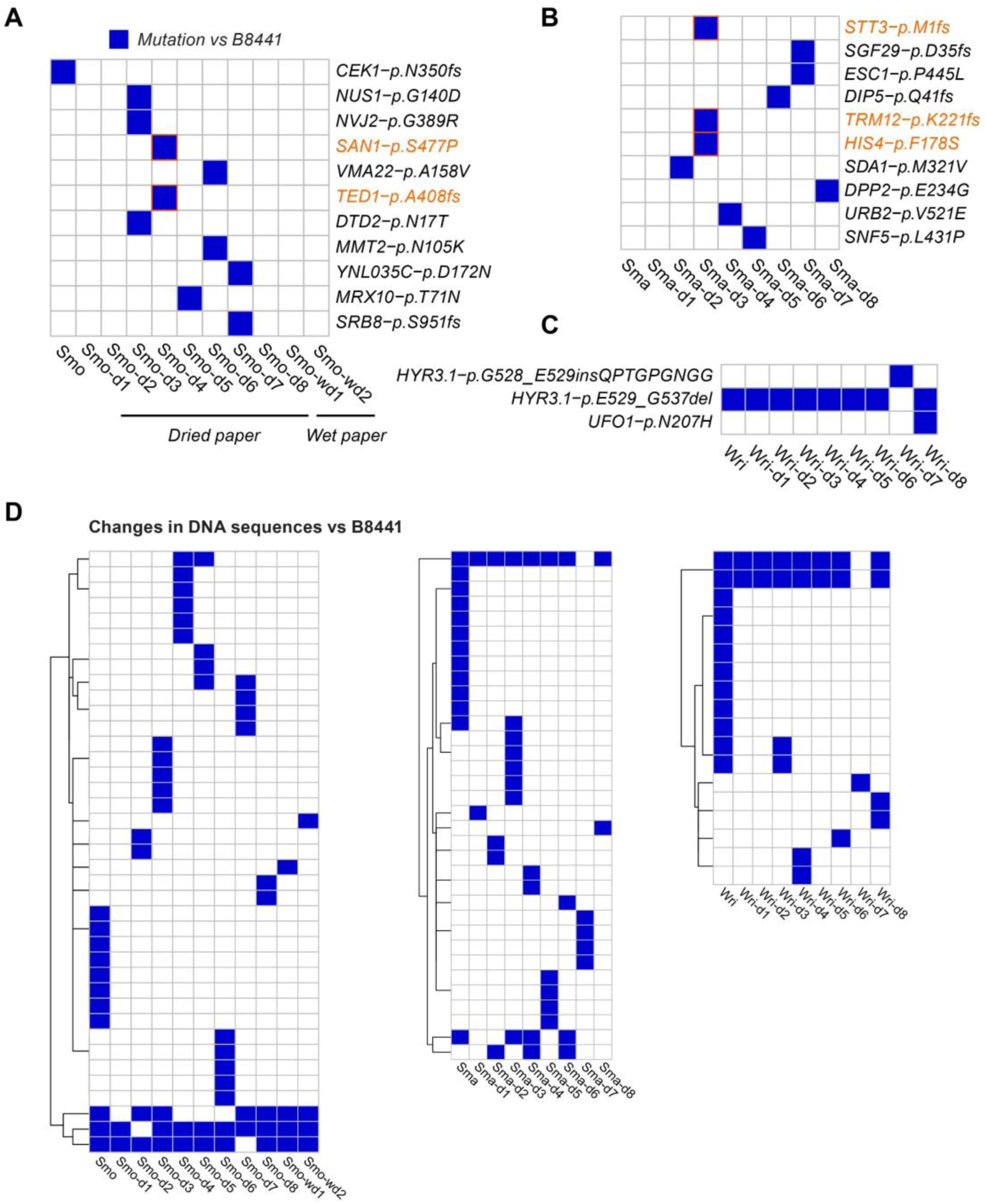
Whole genome sequencing (WGS) analysis of distinct derivative clones reconstituted from filter papers. **A-C.** Heatmaps show non-synonymous and frameshift mutations in coding genes of original strains and their derivative clones. Orange color highlights potential mechanisms of voriconazole susceptibility. **D.** Mutations in DNA sequences of derivatives clones compared to their parental strains.

Although some derivatives of the Smo morphotype exhibited altered susceptibility to VRC and 5FC, mutations in genes associated with antifungal resistance traits were not observed. However, non-synonymous mutations in the ubiquitin-protein ligase San1 (B9J08_003302 – p.S477P) and an ER to Golgi vesicle-mediated transporter Ted1 (B9J08_004004 – p.A408fs) may account for the increased VRC resistance of Smo-d3, while mutations in the oligosaccharyl transferase *STT3* (B9J08_000148 - p.M1fs), the S-adenosylmethionine-dependent methyltransferase *TRM12* (p.K221fs), and a histidinol dehydrogenase *HIS4* (p.F178S) may account for the VRC hypersensitivity of Sma-d3. The molecular functions of those genes and their mode of involvement need to be addressed in the future. Smo-d2 and Smo-d3 displayed increased susceptibility to 5FC; Smo-d2 did not carry non-synonymous mutations, but several mutations were present in non-coding regions. These may involve non-coding RNAs, promoter, or terminator sequences affecting gene expression, thereby contributing to altered drug susceptibility **(Fig. 4D)**. Taken together, our data suggest that *C. auris* pan-antifungal traits are resulting from combinatorial mechanisms, which all together lead to a compound resistance phenotype, including stochastic mutagenic events in the absence of selective pressure.

## DISCUSSION

We here show that storing or preserving *C. auris* strains on dried filter paper may promote the emergence of new morphotypes with altered fitness and drug resistance traits upon reconstitution of pure isolates. This effect may be mitigated by maintaining humidity during storage to reduce osmotic and/or drought stress. Remarkably, our data support the previous notion that extreme drought stress paired with increased salinity *C. auris* encounters in drying environmental wetlands has constituted a major driving force of mutational adaptation, since it equips the pathogen with thermos- and halotolerance, antifungal resistance, as well as the ability to expand its host range [33]. Importantly, the data also highlights a potential and hitherto unrecognized pitfall in routine laboratory strain exchanges. The data also raises serious concerns as to how rapid *C. auris* can evolve in healthcare facilities or when colonizing individuals. Strikingly, the persistence of *C. auris* on abiotic surfaces, combined with host stressors, may be sufficient to trigger random mutational adaptations including morphogenetic, changes in the absence of antifungal selection pressure or other *in vivo* challenges in the host settings. This intrinsic genetic makeup of *C. auris* can allow for the emergence of new pathogenic traits and efficient anti-fungal resistance.

While we cannot entirely exclude that the original patient isolate resulted from a mixed infection, we show that mutations appear when drying isolates on sterile filter paper. Indeed, our experimental data demonstrate that random *de novo* mutations emerge after two weeks of storage at room temperature, even under humidity conditions. These mutations occur in previously unidentified resistance-related genes including non-coding regions [47], leading to altered antifungal susceptibility in several derivative clones. This finding suggests that *C. auris* benefits from a high mutation rate, which could drive antifungal resistance even in the absence of drug exposure. Importantly, our results highlight the need to re-evaluate and improve current protocols for strain transfer between research laboratories, calling for standard operating procedures when exchanging strains between groups.

Although *C. auris* does not exhibit hyphal growth as prominently as *Candida albicans*, hyphae formation occurs in multiple clinical isolates across all major clades under specific culture conditions or in infection models [48–50]. This filamentation is likely an epigenetic phenomenon [48,51], influenced by environmental triggers such as osmotic stress or DNA damage [52,53]. Our data suggest that mutations affecting cell separation, growth and DNA damage response **(Fig. 2A, B)**, which may establish a genetic makeup with a higher propensity for epigenetic switching as observed in Wri morphotypes **(Fig. 3A, B)**. Although filamentous growth in *C. auris* appears limited, the data indicate that the combination of a high mutation rate and epigenetic regulation may enhance virulence traits in certain stress-rich environments. All in all, this enhances the pathogen adaptability related to both virulence and drug resistance.

Regarding drug resistance, the WGS analysis identified novel mutations in the transcription factors Tac1b and Mrr1, both of which are master regulators of the ABC efflux transporters Cdr1 and the major facilitator permease Mdr1, respectively, the main mediators of azole resistance in *Candida* spp [12–14,40,42]. Additionally, we discover a novel loss-of-function mutation in *FKS2*, leading to increased susceptibility to caspofungin owing to an inactive truncated Fks2 glucan synthase. These findings highlight the potential of Fks2 for mediating caspofungin resistance in *C. auris*, a function that has previously been underexplored. Indeed, although Fks1 is the primary glucan synthase with high expression *in vitro* in *C. glabrata*, recent findings reveal that *FKS2* is expressed at equivalent or even higher levels than *FKS1* during stationary phase or when facing host defense [54]. Our data underscore the pivotal relevance of Fks2, Tac1b, and Mrr1 in antifungal resistance and suggests a need for deep mutational scanning of these genes in primary clinical isolates. Such efforts would contribute to the development of a comprehensive reference database for antifungal resistance prediction and diagnostics for *C. auris* and potentially other *Candida* pathogens.

Of note, we also observe a notable increase in 5FC susceptibility among several Smo derivative clones. However, this effect is unlikely to promote fungicidal effect, it rather reflects 5FC-intrinsic metabolic effects. Although these strains exhibited growth inhibition, the response was gradual and extended across a broad concentration range. From a clinical perspective, 5FC treatment failures in *Candida* spp are typically linked to acquired mutations in the *FCY1–FUR1–FCY2* pathway [8,18,55]. The altered susceptibilities observed in our clones may result from mutations affecting metabolic pathways, potentially leading to fitness alterations **(Figure 3C)**.

Moreover, azole susceptibilities are significantly altered in several derivative clones (Smo-d3, Sma-d3); however, no clear mutations were detected in well-characterized resistance genes. This is not unexpected, as azole resistance mechanisms in *C. auris* are complex, highly redundant and multifactorial. Thus, azole resistance is not solely associated with mutations in the ergosterol biosynthesis pathway (e.g., *ERG11*), but also with the overexpression of drug efflux transporters including *MDR1*, *CDR1*, other membrane transporters as well as the putative phospholipid exchange factor *PDR16* [11,12,14,17]. These membrane transporters are regulated by multiple and specific, partially overlapping transcriptional networks, involving *Tac1b*, *Mrr1*, *Upc2*, and *Rpn4* [14,40,56,57]. Therefore, the rapid accumulation of mutations driving antifungal resistance after only a few cell divisions represents a remarkable trait of *C. auris*.

In summary, we show that drying of an exemplary *C. auris* clinical isolate on filter paper promotes rapid mutagenic events that promote altered morphogenesis and drug resistance upon reconstitution. A similar phenomenon may also occur in other strains [58], as several studies have reported mixed populations within individual clinical isolates or diversification within a single strain driven by both genetic mutations and epigenetic regulation [4,32]. Thus, we propose that the community establishes standards for interlaboratory exchange of isolates as well as phenotypic protocols for verifying phenotypes of reconstituted strains.

## METHODS

### Media, culture and morphology examination

*C. auris* strains preserved on filter paper received from collaborating laboratories were reconstituted by placing onto YPD agar plates and incubating at 37 °C. Three distinct colony morphotypes emerging were designated Smooth (Smo), Small (Sma), and Filament (Fil). Colonies of pure morphotypes were restreaked on YPD agar, followed by conservation as frozen stock at −80 °C in 20% glycerol. Three distinct colony morphotypes, designated Smooth, Small, and Filament, were identified, isolated, and stored at −80 °C in 20% glycerol stocks. Each morphotype was cultured overnight at 37 °C in YPD broth. Cultures were washed twice with sterile distilled water and adjusted to a final concentration of 10^8^ cells/ml. A 50 µL aliquot of the suspension was spotted onto 2Dcm^2^ sterile filter papers and allowed to air dry. Dried filter papers were wrapped in sterile aluminum foil for storage. As a control, moist filter papers were placed into sterile microcentrifuge tubes sealed with paraffin film. Both test and control samples were stored at room temperature for two weeks. After storage, filter papers were submerged in 1 mL of sterile distilled water, vortexed briefly, serially diluted, and plated onto YPD agar. Colonies were incubated at 37D°C and assessed morphological characteristics. Eight colonies exhibiting distinct morphologies were isolated. Among them, isolate d1 retained morphology similar to the original strains, while others displayed altered phenotypes. Unless otherwise stated, *Candida* strains were routinely cultured overnight in YPD medium at 37 °C for all subsequent experiments.

### Antifungal susceptibility testing

Antifungal susceptibility was assessed using a dose–response assay following the Clinical and Laboratory Standards Institute (CLSI) guidelines [32,59]. The assay was performed in RPMI 1640 medium supplemented with 35 g/L MOPS and buffered to pHD7.0. Fungal strains were first grown in YPD medium to the exponential phase, then washed and adjusted to the appropriate concentration. Approximately 500-2,500 colony-forming units (CFUs) were inoculated into each well of a 96-well microtiter plate containing antifungal agents in a two-fold serial dilution series. Control wells without antifungal drugs were included. Plates were incubated at 37 °C for 24Dhours, and optical density was measured at 600 nm using a PerkinElmer Victor Nivo Multimode plate reader. Minimum inhibitory concentrations (MICs) were defined as the lowest drug concentration that resulted in ≥50% growth inhibition relative to the no-drug control for all antifungal agents, except for amphotericin B, where a 90% inhibition threshold was used.

### DNA extraction and Sanger sequencing

Genomic DNA was extracted from *Candida* strains grown overnight in 5 mL of YPD medium at 30D°C using the phenol:chloroform:isoamyl alcohol method [21]. Cultures were centrifuged at 3,000Drpm for 3Dminutes, and the resulting cell pellet was resuspended in 200 µL of Yeast Lysis Buffer (composed of 4 mL Triton X-100, 20 mL 10% SDS, 4DmL 5DM NaCl, 400 µL 0.5DM EDTA, and 1DM Tris-HCl, pH 8.0). An equal volume (200 µL) of phenol:chloroform:isoamyl alcohol (25:24:1) and 200 mg of acid-washed glass beads (Sigma) were added to the suspension. Cell lysis was performed using a FastPrep homogenizer for 45Dseconds at 6Dm/s, repeated twice. After lysis, 200 µL of TE buffer (pH 8.0) was added, and samples were centrifuged at 13,000×g for 5Dminutes. The aqueous phase was carefully transferred to a new microcentrifuge tube.

DNA was precipitated by adding 1 mL of 100% ethanol and incubating the mixture at −20 °C for 20Dminutes. Following centrifugation at 4 °C for 10Dminutes, the supernatant was discarded, and the DNA pellet was resuspended in 400 µL TE buffer containing 4 µL of RNase A (10 mg/mL). Samples were vortexed and incubated at 37 °C for 20Dminutes. For further purification, 10 µL of 4DM ammonium acetate and 1 mL of 100% ethanol were added, followed by a second incubation at −20 °C for 20Dminutes. Samples were centrifuged at 4 °C for 10Dminutes, the supernatant was removed, and the pellet was washed with 1 mL of 70% ethanol. After air-drying for 5Dminutes, the DNA pellet was resuspended in 50 µL of TE buffer or sterile water and incubated at 37 °C for 5Dminutes before downstream applications. A fragment of the *FKS2* gene was amplified from genomic DNA using the primer pair: Forward – 5′-CTCAAGCAACACTCACCGTC-3′; Reverse – 5′-AAAGACGTTTCTCCAAGGCG-3′. PCR products were purified with GeneJET PCR Purification Kit (ThermoScientific, K0702) and submitted for Sanger sequencing using the Mix2Seq Kit NightXpress (Eurofins Genomics) according to the manufacturer’s instructions.

### Whole-genome sequencing and bioinformatics analyses

The quality of extracted genomic DNA was assessed using a Qubit 3.0 fluorometer and agarose gel electrophoresis. DNA libraries were prepared using the VAHTS Universal Plus DNA Library Prep Kit for Illumina (ND617). Fragment size distribution and overall library quality were evaluated using the Qsep-400 capillary electrophoresis system. Sequencing was carried out on the Illumina NovaSeq platform in paired-end 150 bp (PE150) mode by Biomarker Technologies (BMK) GmbH, Germany.

Whole genome sequencing (WGS) data was analyzed using the CDC’s MycoSNPs pipeline [60]. Variants were filtered based on the following criteria: QD < 2.0, FS > 60.0, MQ < 40.0, or DP < 10. To reduce false positives and increase variant confidence, genotype masking was applied using thresholds of genotype quality ≥50, read depth ≥10, and alternate allele frequency ≥80% in successive filtering steps [8].

### Data availability and statistical methods

WGS data will be deposited via the NCBI Sequence Read Archive (SRA) BioProject PRJNA1311805. The entire bioinformatic workflow adapted from CDC mycoSNP [60] as previously published via Github at https://github.com/kakulab/5FC-Evo-2024 [8].

## Disclaimer

The funding sources had no role in the design, execution, or reporting of this study.

## Acknowledgements

We express our gratitude to all KaKulab members for their technical support and experimental advice.

## Author contributions

Conceptualization: TPC, KK; Methodology: TPC, PLL, NC, KK; Investigation: TPC, D-M. L-M, AC, MC, NC; Formal analysis: TPC, D-M. L-M; Visualization: TPC; Funding acquisition: TPC, KK; Project administration: TPC, KK; Supervision: KK; Writing – original draft: TPC, KK; Writing – review & editing: all authors.

---

**Figure S1.**
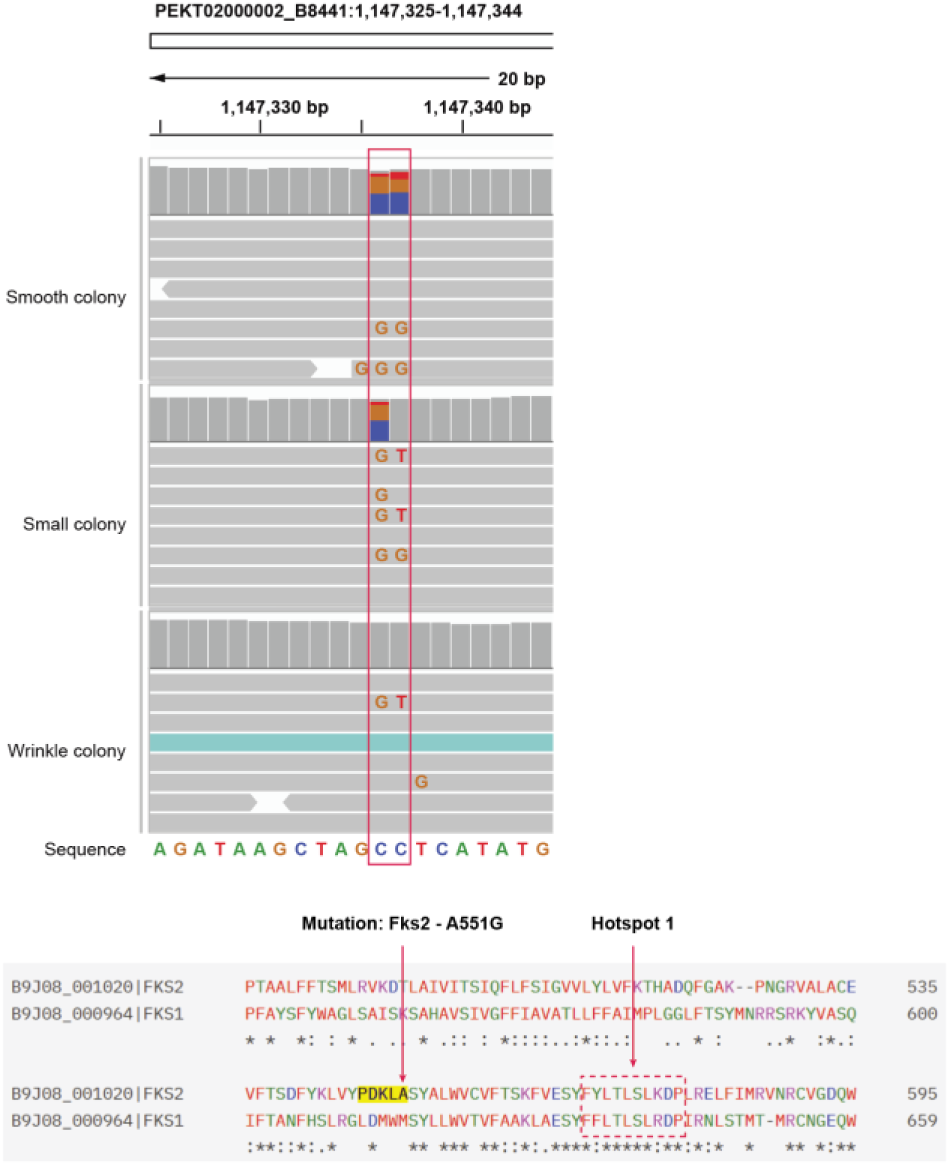
A new mutation in *FKS2* is related to caspofungin resistance from WGS datasets.

